# Mitochondrial dysfunction in skeletal muscle of fukutin deficient mice is resistant to exercise- and AICAR-induced rescue

**DOI:** 10.1101/2020.05.27.118844

**Authors:** W. Michael Southern, Anna S. Nichenko, Anita E. Qualls, Kensey Portman, Ariel Gidon, Aaron Beedle, Jarrod A. Call

**Affiliations:** Department of Kinesiology, University of Georgia, Athens, GA 30602, USA; Regenerative Bioscience Center, University of Georgia, Athens, GA 30602, USA; Department of Pharmaceutical Sciences, SUNY at Binghamton, Binghamton NY 13902, USA; Department of Pharmaceutical and Biomedical Sciences, University of Georgia, Athens, GA 30602 USA

**Keywords:** muscle torque, mitochondrial respiration, mitochondrial oxygen consumption, muscle regeneration, oxidative plasticity, dystroglycan, fukutin, muscular dystrophy

## Abstract

Disruptions in the dystrophin-glycoprotein complex (DGC) are clearly the primary basis underlying various forms of muscular dystrophies and dystroglycanopathies, but the cellular consequences of DGC disruption are still being investigated. Mitochondrial abnormalities are becoming an apparent consequence and contributor to dystrophy disease pathology. Herein, we demonstrate that muscle-specific deletion of the fukutin gene [Myf5/*fktn-*KO mice (KO)], a model of secondary dystroglycanopathy, results in ~30% lower muscle strength (P<0.001) and 16% lower mitochondrial function (P=0.002) compared to healthy littermate controls (LM). We also observed ~80% lower PGC-1α signaling (P=0.004), a primary transcription factor for mitochondrial biogenesis, in KO mice that likely contributes to the mitochondrial defects. PGC-1α is post-translationally regulated via phosphorylation by AMPK. Treatment with the AMPK agonist AICAR (5-aminoimidazole-4-carboxamide ribonucleotide) failed to rescue mitochondrial deficits in KO mice (P=0.458) but did have beneficial (~30% greater) effects on recovery of muscle contractility following injury in both LM and KO mice compared to saline treatment (P=0.006). The beneficial effects of AMPK stimulation via AICAR on muscle function may be partially explained by AMPK’s other role of regulating skeletal muscle autophagy, a cellular process critical for clearance of damaged and/or dysfunctional organelles. Two primary conclusions can be drawn from this data, 1) fukutin deletion produces intrinsic muscular metabolic defects that likely contribute to dystroglycanopathy disease pathology, and 2) AICAR treatment accelerates recovery of muscle function following injury suggesting AMPK signaling as a possible target for therapeutic strategies.

## Introduction

Muscular dystrophy is characterized by progressive muscle weakness and atrophy due to the loss of structural integrity of the dystrophin-glycoprotein complex (DGC) within muscle cells. The DGC provides a critical link between the extracellular matrix of skeletal muscle cells with the intracellular cytoskeleton^1,2^. Without proper function of this protein complex, muscle fibers are susceptible to injury, leading to a progressive muscle dysfunction^3^. One important functional component of the DGC is the extracellular protein α-dystroglycan (αDG) that acts as a receptor for extracellular matrix proteins such as laminin^4^. Disruptions in the function of αDG give rise to dystroglycanopathies characterized by early onset of muscle weakness and progressive loss of muscle function^5^. The fukutin protein, encoded by the *FKTN* gene, is one of a cascade of proteins responsible for glycosylation of αDG, a post-translational modification necessary for αDG binding to laminin and other extracellular matrix receptors^6^. Mutations in the *FKTN* gene have been directly linked to αDG dysfunction and subsequent muscle dystroglycanopathy^7^. A fukutin knockout mouse (KO) model accurately recapitulates many of the characteristics observed in human dystroglycanopathies as the muscles are characterized by chronic weakness, a perpetual cycle of injury and repair, and abnormal muscle regeneration^8,9^. That said, several pathological features of dystroglycanopathies remain undefined, and herein we sought to interrogate mitochondrial function based on our previous work which eluded to possible mitochondrial alterations in fukutin KO muscle^10^.

Skeletal muscle mitochondria are a critical network of organelles within the muscle cell responsible for energy production, redox control, ion homeostasis, and signaling. The health of the mitochondrial network can influence the health and function of the muscle cell. Maintaining a healthy mitochondrial network is a balance between the addition of new mitochondria to the network, i.e. mitochondrial biogenesis, and the removal of damaged or dysfunctional mitochondria from the network, i.e. mitochondrial autophagy. Previously, we identified a mild reduction in succinate dehydrogenase activity, a marker of mitochondrial content, in fukutin KO mice compared to wildtype controls suggesting that mitochondrial maintenance (biogenesis and/or autophagy) is detrimentally impacted by fukutin KO-induced disruption of the DGC^10^. Our findings are in line with other studies highlighting compromised mitochondrial function and content in mouse models of DGC, suggesting that DGC disruptions are linked to the metabolic health of the muscle.

If mitochondrial maintenance is altered in fukutin KO mice, then targeting mitochondria biogenesis or autophagy may be a viable therapy to explore for treatment of secondary dystroglycanopathies. One potential target is the cellular energy-sensing AMP-activated protein kinase, AMPK, that is critical for post-translational modifications to *PGC-1α* (a metabolic gene transcription factor) and ULK1 (a kinase that initiates autophagosome assembly)^11,12^. As demonstrated by Pauly et al., pharmacological treatment of dystrophin-deficient *mdx* mice with the AMPK agonist AICAR enhanced autophagy and ameliorated several of the pathological characteristics common to *mdx* mice including muscle weakness and some metabolic defects^13^. These results highlight the therapeutic potential of AMPK-stimulation via AICAR to potentially correct metabolic and contractile abnormalities in fukutin KO mice.

The purpose of this study was to determine the extent to which fukutin KO disrupted mitochondrial function in skeletal muscle and then test if enhancing mitochondrial quality in KO mice affects muscle function. We hypothesized that fukutin KO mice will have lower mitochondrial function than control mice, and that enhancing the mitochondrial network will be beneficial for muscle function. To test this hypothesis, we utilized highly sensitive and reproducible electrophysiological techniques to measure both muscle contractility and mitochondrial quality in KO mice.

## Methods

### Ethics Statement

#### Approval

All animal protocols were approved by the University of Georgia Animal Care and Use Committee under the national guidelines set by the Association for Assessment and Accreditation of Laboratory Animal Care.

### Animals and Study Design

Male and female Myf5/*Fktn* knockout (KO) mice (previously described^9^) and littermate controls (LM) 8-12 weeks of age were maintained at 20-23° C on a 12:12 hour light:dark cycle with food and water *ad libitum*. LM and KO mice (63% male, 37% female) were randomized into either saline (LM n= 13, KO n= 9) or AICAR (LM n= 13, KO n=7) treatment group. Mice received daily injections of saline (200 μl) or AICAR (500mg/kg BM) for a total of 4 weeks^11,13^. After 2 weeks of treatment, the left hindlimbs (anterior and posterior compartments) of all mice were injured with 40 μl of cardiotoxin (Naja kaouthia, 0.071 mg/ml; catalog no. C9759, Sigma-Aldrich); the contralateral limb served as the uninjured control. Following the muscle injury, all mice continued to receive treatment for 2 weeks at which point they were sacrificed. Immediately prior to sacrifice, *in vivo* peak isometric torque was assessed on the dorsiflexor muscles of the left (injured) mouse hindlimb. At sacrifice, injured and uninjured gastrocnemius and tibialis anterior muscles were extracted and weighed. A fraction of the gastrocnemius was immediately processed for measurements of mitochondrial function, while the remaining portion was homogenized and prepared for assessment of mitochondrial enzyme activities and western blotting. The tibialis anterior muscle was processed for histology and microscopy and stored at −80°C.

### *In Vivo* Muscle Function

*In vivo* peak isometric torque of the ankle dorsiflexors was assessed as previously described^14^. Briefly, anesthesia was induced using an induction chamber and 5% isoflurane in oxygen. Anesthesia was maintained using 1.5% isoflurane at an oxygen flow rate of 0.4 L/min. The left hindlimb was depilated and aseptically prepared and the foot placed in a foot-plate attached to a servomotor (Model 300C-LR; Aurora Scientific, Aurora, Ontario, Canada). Platinum-iridium needles (Model E2-12; Grass Technologies, West Warwick, RI) were placed on either side of the left common peroneal nerve to elicit contraction of the dorsi-flexors muscles. Peak isometric torque was defined as the greatest torque measured by a 300B-LR servomotor (Aurora Scientific, Aurora, Ontario, Canada) during a 200-ms stimulation using 1-ms square-wave pulses at 300 Hz and increasing voltage from 3.0 to 9.0 V (models S48 and SIU5; Grass Technologies). All torque values were normalized by mouse body mass.

### Acute stimulation protocol

To assess the integrity of mitochondrial biogenesis signaling in KO muscle, 8 additional untreated mice (n = 4 LM; n = 4 KO) underwent a 30 minute in vivo electrical stimulation protocol to simulate an acute bout of exercise as previously described^15^. Briefly, mice were anesthetized with isoflurane (1.5–2.0%) and platinum-iridium needle electrodes were placed around the sciatic nerve. The electrical stimulation protocol consisted of 10 sets of 1800 contractions (parameters: pulse frequency = 100, pulse width = 0.1, pulses per train = 1, train frequency = 10 Hz) conducted over 30 minutes. At the end of the protocol, gastrocnemius muscles were quickly harvested, flash frozen in liquid nitrogen, and stored at −80 °C for qRT-PCR analysis.

### Mitochondrial Assays

Mitochondrial function was assessed as previously described^15^. Briefly, portions of the medial and lateral gastrocnemius muscles from uninjured limbs were dissected on a chilled aluminum block in 4°C buffer X (7.23mM K_2_EGTA, 2.77mM Ca K_2_EGTA, 20mM imidazole, 20mM taurine, 5.7mM ATP, 14.3mM PCr, 6.56mM MgCl_2_-6H_2_O, 50mM k-MES) into thin muscle fiber bundles (<1mg). Muscle fibers were permeabilized in buffer X and saponin (50 μg/ml) for 30 minutes at 4°C. Muscle fiber bundles were rinsed for 15 minutes in buffer Z (105mM k-MES, 30mM KCl, 10mM KH_2_PO_4_, 5mM MgCl_2_, 0.5 mg/ml BSA, 1mM EGTA) at 4°C. All measurements were performed using a Clark-type electrode (Oxygraph Plus System, Hansatech Instruments, UK) at 25° C. Muscle fiber bundles (~2.5 mg for all samples) were added to the chamber and mitochondrial leak respiration (addition of 10mM glutamate, 5mM malate) and mitochondrial state 3 respiration (addition of 2.5mM ADP and 10mM succinate) were measured. Cytochrome *c* (10μM) was also added to measure the integrity of the outer mitochondrial membrane. Mitochondrial respiration was terminated by the addition of cyanide (2 mM).

Mitochondrial content was assessed using citrate synthase and succinate dehydrogenase enzyme activities as previously described^16,17^. Briefly, a fraction of the gastrocnemius muscles were homogenized in a glass tissue grinder in 33 mM phosphate buffer (pH = 7.0) at a muscle:buffer ratio of 1:20. Citrate synthase activity was assessed by combining 5 μl of homogenate, 173.74 μl of 100 mM Tris buffer (pH 8.0), 17.51 μl of DTNB, 8.75 μl of acetyl CoA, and 20 μl of oxaloacetate in a well of a 96 well plate and measuring absorbance at 405 nm every 10 seconds for 3 minutes. Succinate dehydrogenase activity was assessed by combining 4 μl of homogenate, 10 μl of 0.5 M sodium succinate, 10 μl of 0.01 M sodium cyanide, and 280 μl of cytochrome c solution (33% 0.1 mM cytochrome c, 13% 0.004 M AlCl_3_/CaCl_2_) in a well of a 96 well plate and measuring absorbance at 550 nm every 20 seconds for 5 minutes. Enzyme activities were normalized to mg of tissue in the sample homogenate.

### Western Blotting

For protein content analysis, 30 μg of total protein from injured and uninjured gastrocnemius muscles were separated by SDS-PAGE, transferred onto a PVDF membrane, and immunoblotted as previously described^16^. The following antibodies (Cell Signaling, Danvers, MA) were used: AMPK (RRID: AB_330331, 1:1000), pAMPK (RRID: AB_330330, 1:1000), Beclin-1 (RRID:AB_1903911, 1:1000), LC3B (RRID:AB_915950, 1:1000), SQSTM1/p62 (RRID: AB_1841064, 1:1000), Bnip3 (RRID:AB_2259284, 1:1000), Drp1 (D6C7, 1:1000). Immunoblots were normalized to total protein in lane and quantified using Bio-Rad Laboratories Image Lab software (Hercules, CA).

### Immunofluorescence Microscopy

Seven-micron tissue cryosections were mounted on microscope slides and processed for immunofluorescence analysis of centrally located nuclei (CNF) and embryonic myosin heavy chain (eMHC) as similar to previous work^8^. Sections were wet with Dulbecco’s PBS (no Ca^2+^ or Mg^2+^, Corning); incubated for 30 min in blocking solution (5% donkey serum [Jackson ImmunoResearch] in dPBS); incubated overnight at 4°C with primary antibody; washed three times; incubated for 30 min with secondary antibody, stains or dyes; washed three times; and mounted with PermaFluor (Thermo Scientific). Reagents used for visualization were: F1.652 against embryonic myosin heavy chain (1:40, DSHB); goat anti-mouse IgG1-Alexa546 (1:500, Invitrogen); membrane marker WGA-coupled Alexa488 (company, 1:100); muscle marker Actin-Alexa647 (company, 1:40); and nuclear stain DAPI (company, 1:10,000). Four filter epifluorescent images were obtained for whole TA sections using the VS-120 slide scanner (Olympus) with 10X objective and imported into ImagePro Premier 3D v9.1 (Media Cybernetics) as 3 color images for analysis (centrally nucleated and eMHC counts: eMHC/WGA-A488/DAPI; fiber size: WGA-A488/Actin A647/DAPI). Centrally nucleated - (CNF) and eMHC-positive fibers were analyzed by manual count out of total fibers for entire TA sections. The smart segmentation function in ImagePro Premier was used to partially automate detection of fiber perimeters. All sections were manually verified for fiber detection and corrected as necessary. Minimum diameter (μm) was collected for all fiber annotations on a section. A bin size of 3.5 μm for fiber sizes was selected by taking the average *h* calculated for each sample using the formula 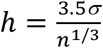 (h = optimized in size, *σ* = standard deviation, n= number of observations) and rounding up to the nearest 0.5. Fiber sizes for each sample were binned and reported as average ± standard error for each group. All image analyses were performed blinded.

### Gene Expression

cDNA was generated from isolated gastrocnemius muscle RNA using a High Capacity cDNA Reverse Transcription Kit (Applied Biosystems, Foster City, CA). iQ SYBR Green Supermix (Bio-Rad) and sequence-specific primers were used to assess mRNA levels for the following genes, *PGC-1α*, (For: 5’-AGC CGT GAC CAC TGA CAA CGA G-3’; Rev: 5’-GCT GCA TGG TTC TGA GTG CTA AG-3’). Gene expression analysis was carried out by qRT-PCR on a Bio-Rad CFX instrument as previously described^15^. Data were normalized to 18S (For: 5’-TTG ATT AAG TCC CTG CCC TTT GT-3’; Rev: 5’-CGA TCC GAG GGC CTA ACT A-3’) and relative gene expression was calculated using the 2^−ΔΔCT^ method.

### Statistics

Data are presented in the results as mean ± SD. A multi-factor repeated measures analysis of variance (ANOVA) was used to analyze genotype (LM, KO), treatment (saline, AICAR), and limb (injured, uninjured) conditions. The between-subject factors were genotype and treatment while the repeated measure factor was limb. Chi-square tests were used to analyze the frequency distribution of fiber cross-sectional areas. Two-way ANOVA was used to analyze the remaining data. All data were required to pass normality (Shapiro-Wilk) and equal variance tests (Brown-Forsythe *F* test) before proceeding with the ANOVA. Differences among groups are only reported where significant interactions were observed and subsequently tested with Tukey’s *post hoc* test using JMP statistical software (SAS, Cary, NC). Group main effects are reported where significant interactions were not observed. An α level of 0.05 was used for all analyses.

## Results

### Characterization of contractile and metabolic function of KO mice

In agreement with previous reports, KO mice had ~21% lower body mass compared to healthy LM control mice (P<0.001; Fig. 1a; Beedle et al, 2012). *In vivo* assessment of peak-isometric torque revealed that KO mice were ~30% weaker than LM controls even after accounting for the differences in body mass (P<0.001; Fig. 1b). State III mitochondrial respiration from permeabilized muscle fibers was also ~16% less in KO mice compared to LM mice suggesting that fukutin deletion may interfere with metabolic processes in skeletal muscle (P=0.002; Fig. 1c).

**Figure 1:**
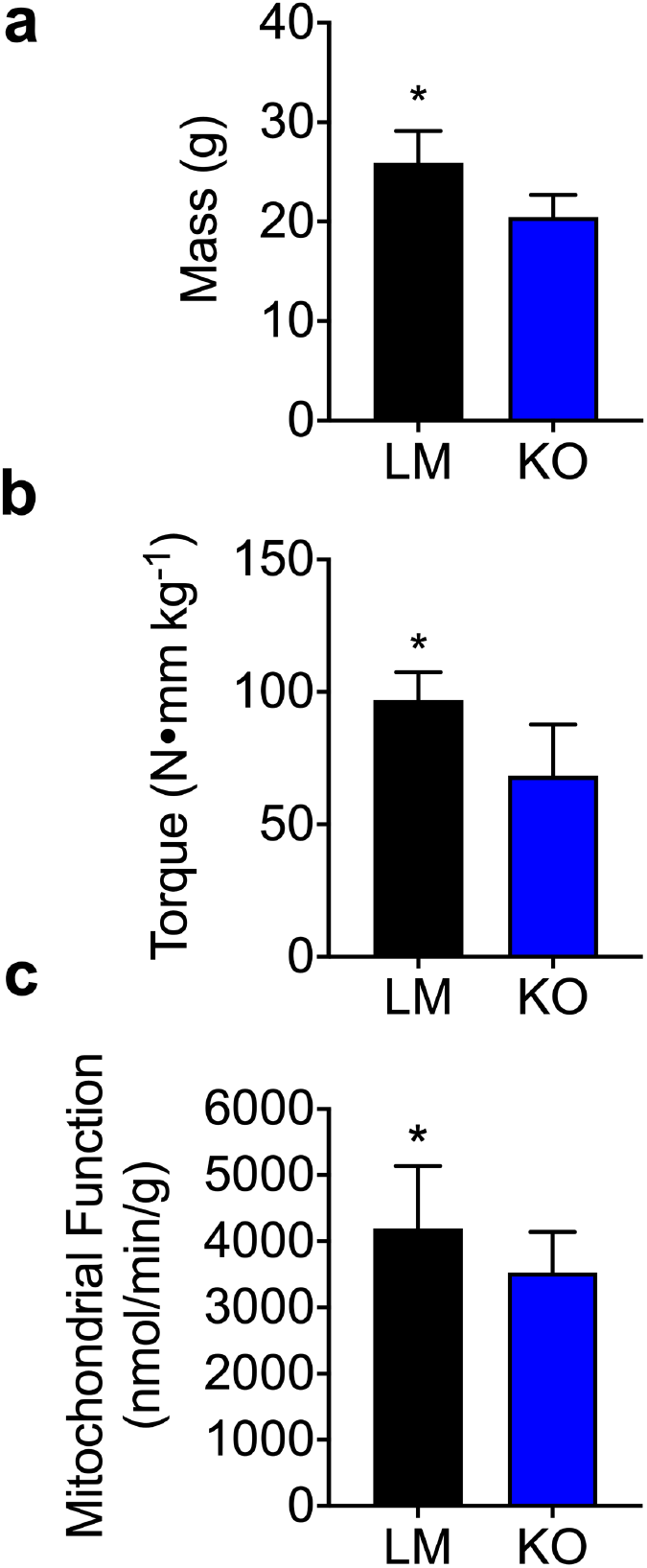
Fukutin KO mice have lower body mass, muscle strength, and mitochondrial function compared to LM control mice. (a) Body mass of LM (n=11) and KO (n=11) mice; *p<0.001.(b) Peak isometric torque of dorsiflexor muscles normalized to body mass of LM (n=13) and KO (n=18) mice; *p<0.001. (c) Mitochondrial function normalized by grams wet weight of permeabilized muscle fibers from LM and KO mice (n ≥ 20 permeabilized fiber bundles from n = 10 mice for each group); *p=0.002. Data are presented as means ± SD.

### Effects of AICAR treatment on mitochondrial content and function in KO mice

In an effort to rescue the loss of mitochondrial function, we treated LM and KO mice with AICAR, a drug previously shown to improve mitochondrial quality in dystrophic muscle via the AMPK-PGC-1α molecular pathway. Interestingly, after 4 weeks of treatment, AICAR did not improve mitochondrial content or function in either LM or KO mice (Fig. 2a-c), but rather resulted in a slight (~7%) reduction in mitochondrial content irrespective of genotype (P=0.010, Fig. 2b-c).

**Figure 2:**
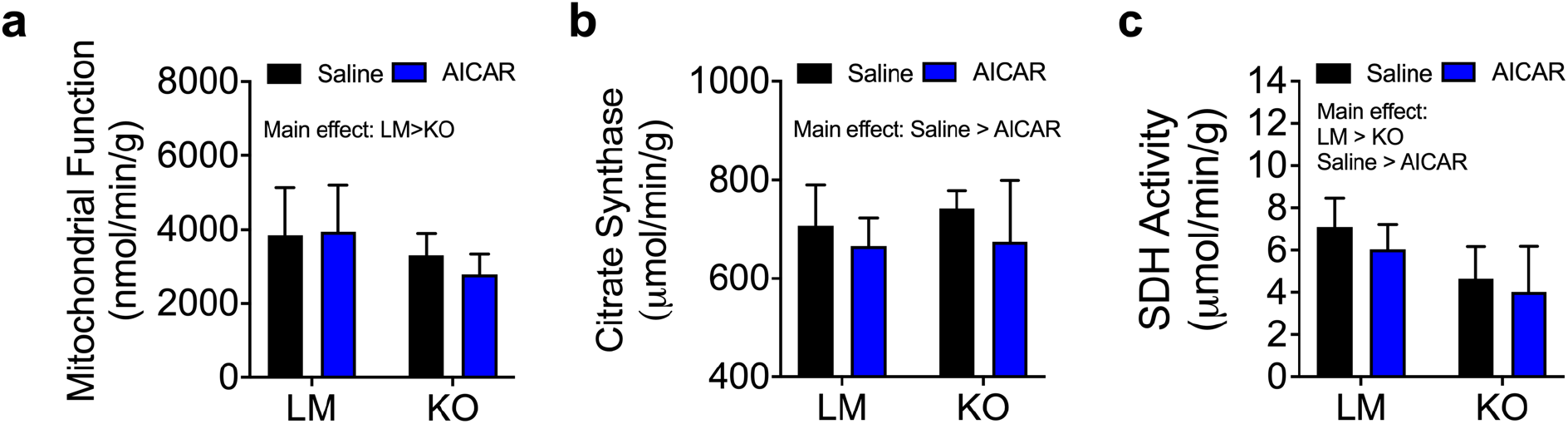
Mitochondrial deficits in KO mice are still present after 4 weeks of AICAR treatment. (a) Mitochondrial function normalized by grams wet weight of permeabilized muscle fibers (main effect of genotype, p=0.004), (b) citrate synthase activity (CS) (two-way ANOVA, p=0.01 for main effect of treatment), and (c) succinate dehydrogenase (SDH) activity (two-way ANOVA, p=0.04 for main effect of treatment and p<0.001 for main effect of genotype) from LM and KO mice after 4 weeks of saline or AICAR treatment (Saline: LM n= 13, KO n= 9; AICAR: LM n= 13, KO n=7). Data are presented as means ± SD.

### Blunted mitochondrial biogenesis in fukutin KO mice

Dysfunctional mitochondrial biogenesis could be a potential mechanism explaining the lower mitochondrial function and content observed in KO mice. In order to test whether KO mice had disrupted mitochondrial biogenesis, we administered an acute exercise stimulus to a hindlimb of LM and KO mice and measured pAMPK content and *PGC-1α* gene expression. Exercised limbs had ~80% greater pAMPK/AMPK protein expression compared to control limbs irrespective of genotype (P=0.040; Fig. 3a-b), demonstrating that exercise-induced AMPK signaling was intact in KO mice. Interestingly, the same did not hold true for post-exercise *PGC-1α* gene expression as LM mice had a ~3-fold increase in *PGC-1α* expression while the KO mice did not have a significant change in expression levels (interaction: P=0.004; Fig. 3c). Together these data suggest that KO mice have blunted mitochondrial biogenesis through impaired activation of *PGC-1α* signaling.

**Figure 3:**
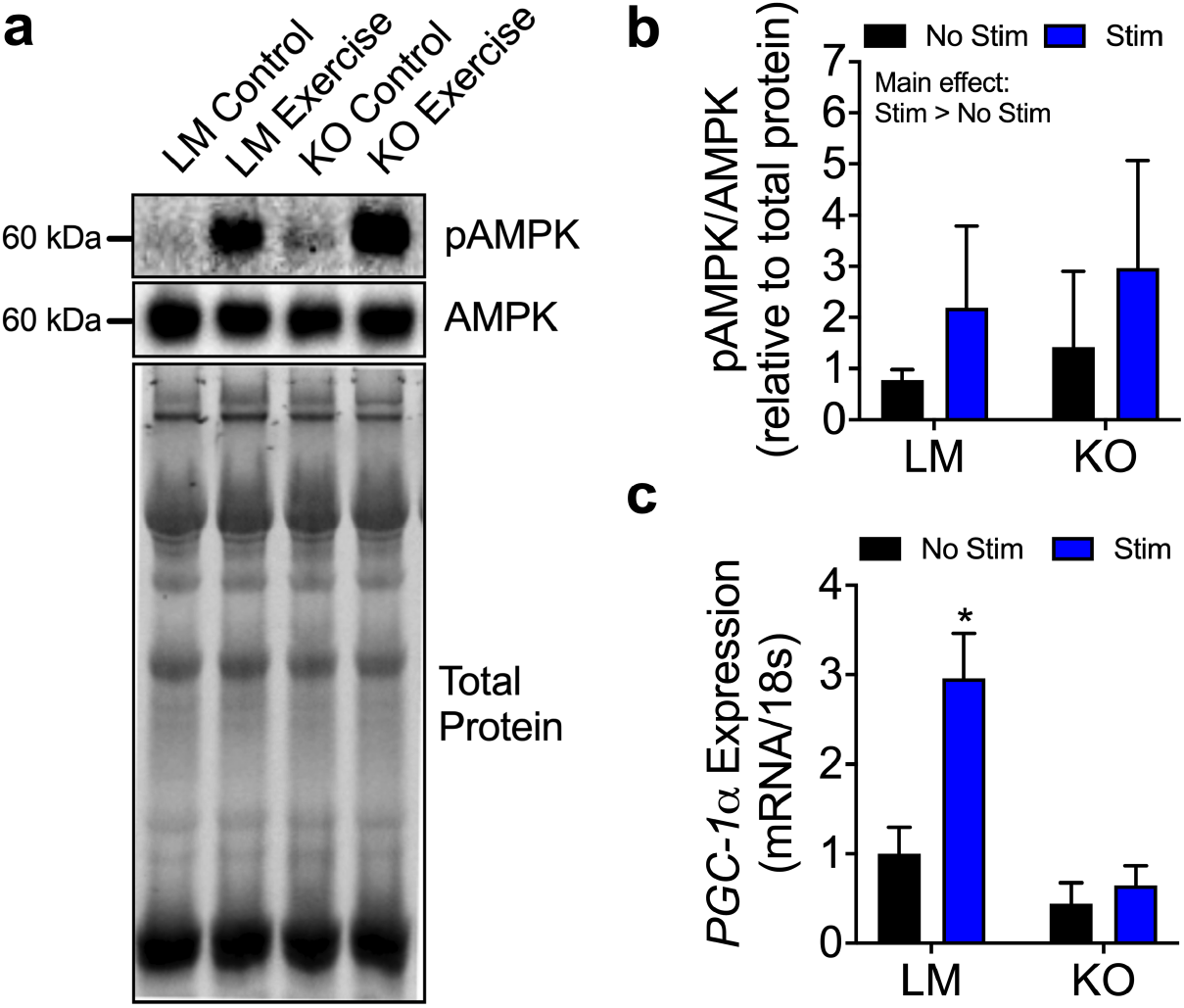
Altered mitochondrial biogenesis signaling in fukutin KO mice. (a) Representative immunoblot and (b) semi-quantitative analysis of pAMPK and AMPK relative protein expression 3 hours after completion of stimulation protocol for stimulated (Stim) and non-stimulated (No Stim) gastrocnemius muscles of LM (n=4) and KO (n=4) mice. Blots are normalized to total protein as a loading control. (c) *PGC-1α* gene expression 3 hours after completion of stimulation protocol for stimulated and non-stimulated limbs of LM and KO mice; two-way repeated measures ANOVA, interaction P < 0.01, * indicates significantly different from all other experimental groups. Data are presented as means ± SD.

### Effects of AICAR treatment on recovery of muscle function following muscle injury

We have previously shown that disruptions in mitochondrial maintenance can have an impact on recovery of muscle contractile function following muscle injury^16^. Thus, in light of the mitochondrial dysfunction in fukutin KO mice, we investigated 1) whether KO mice would have delayed recovery of muscle function following injury and 2) whether AICAR treatment would enhance the recovery process. Prior to injury, KO mice had ~22% lower peak-isometric torque compared to LM controls, independent of saline or AICAR treatment (P<0.001; Fig. 4a). Two weeks after the injury, the untreated LM and KO mice had recovered to ~60% of pre-injury peak-isometric torque indicating that KO mice, despite having mitochondrial abnormalities, did not have delayed recovery of muscle contractility. Interestingly, AICAR-treated mice had ~30% greater peak-isometric torque 14 days after injury compared to untreated mice independent of genotype (P=0.006), which suggests that AICAR treatment reduced damage and/or enhanced the recovery of muscle function for both LM and KO mice (Fig. 4b).

**Figure 4:**
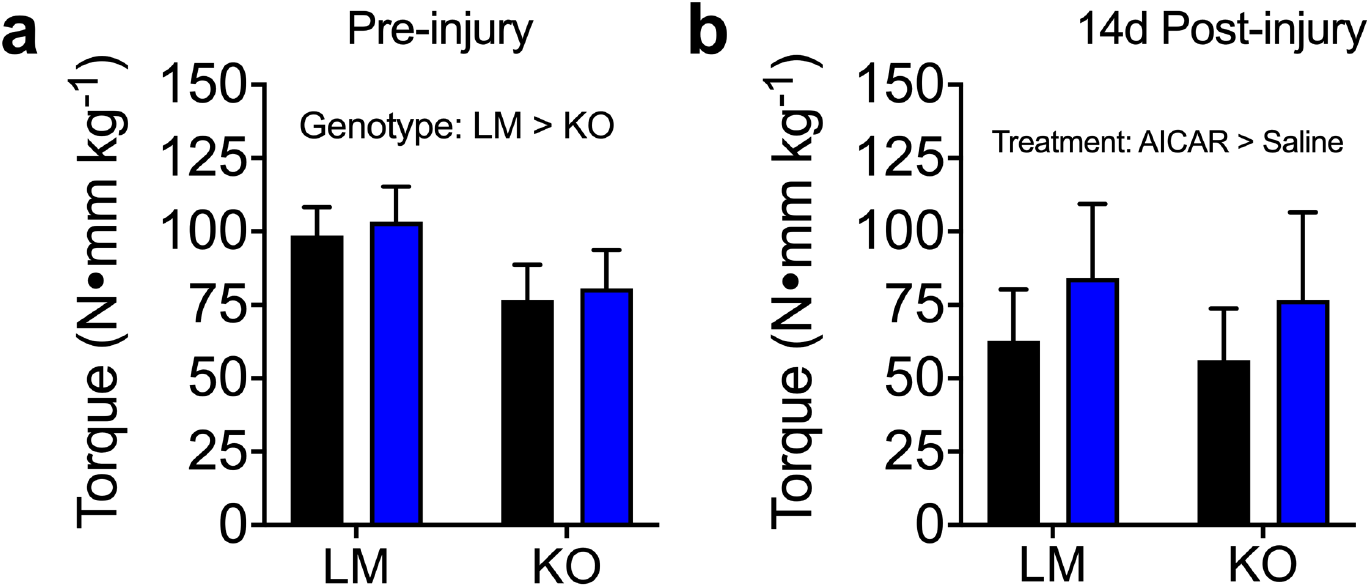
AICAR treatment aids in recovery of muscle strength following muscle injury. (a) Pre-cardiotoxin injury muscle strength from LM and KO mice treated with saline (LM n=13, KO n=9) or AICAR (LM n=13, KO n=7; two-way ANOVA, main effect of genotype P<0.001) (b) Muscle strength 14 days after cardiotoxin injury for LM and KO mice treated with saline (LM n=12, KO n=9) or AICAR (LM n=10, KO n=6; main effect of treatment P=0.006). Data are presented as means ± SD.

Immunohistological analysis of muscle fibers expressing embryonic myosin heavy chain (eMHC) and centrally-nucleated fibers (CNF) was performed to interrogate aspects of the muscle regeneration process in LM control and KO mice. Specifically, eMHC was interpreted as indicating acute/ongoing fiber regeneration and CNF was interpreted as evidence of past/historical fiber regeneration. There was a significant interaction between genotype and limb (uninjured vs. injured) for CNF indicating greater historical regeneration in injured LM muscle and both uninjured and inured KO muscle compared to uninjured LM muscle (P<0.001; Fig. 5a). There was a significant interaction between genotype and treatment (saline vs. AICAR) for eMHC indicating less ongoing regeneration in AICAR-treated KO muscles (P=0.014; Fig. 5b). These data support the notion that muscle damage (CNF) was not affected by AICAR, rather AICAR enhances muscle regeneration after injury (Fig. 5a-b). While all groups had similar muscle fiber numbers (Fig. 5c, Chi-squared analysis of muscle fiber size revealed a distribution shift to the left for the KO mice compared to LM mice and injured muscles compared to uninjured muscles (irrespective of treatment, Fig. 5d-e). Together with the analyses of mitochondrial respiration and AMPK and PGC1α signaling (fig. 2 and 3), these data suggest that AICAR treatment aids in recovery of muscle function, but this action is independent of changes in mitochondrial function.

**Figure 5:**
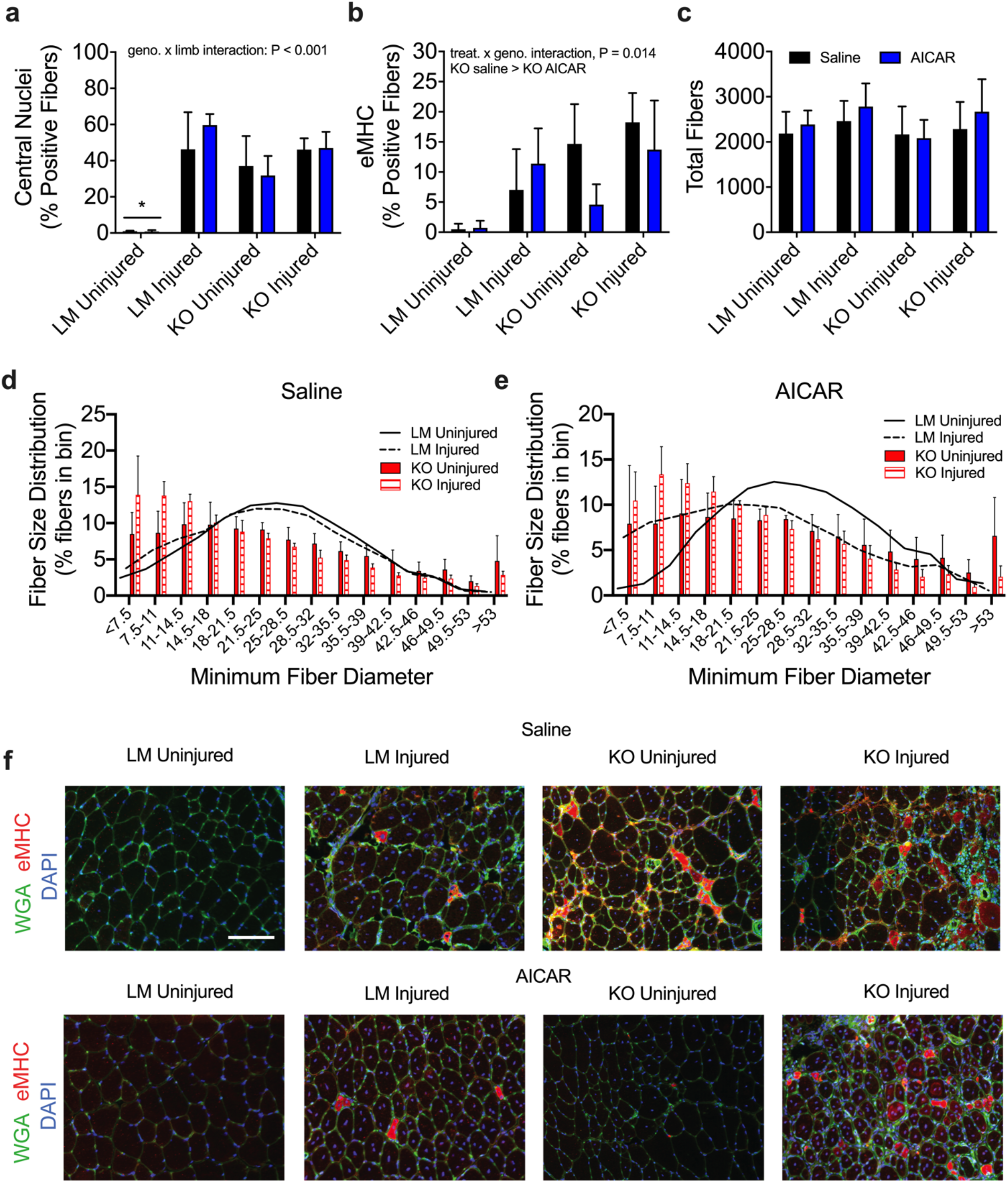
The effects of AICAR treatment on metrics of muscle fiber regeneration for uninjured and injured muscles from LM and KO mice. (a) Percent of total fiber with centrally located nuclei (two-way repeated measures ANOVA, genotype x limb interaction P<0.001) from LM and KO mice treated with saline (LM n=5, KO n=5) or AICAR (LM n=5, KO n=5). (b) Percent of total fibers with positive embryonic myosin heavy chain (eMHC) staining (treatment x genotype interaction, P=0.014). (c) Total number of fibers. (d-e) Fiber size distribution for injured and uninjured muscles from LM (uninjured = solid line, injured = dashed line) and KO (uninjured = solid bar, injured = hatched bar) mice treated with (d) Saline or (e) AICAR. Chi-squared analysis, data are presented as means ± SD. (f) Representative injured and uninjured tibialis anterior images from LM and KO mice treated with saline (top row) or AICAR (bottom row). Wheat germ agglutinin (WGA) (green), embryonic myosin heavy chain (red), and nuclear (blue; DAPI) stains are shown for all groups. Scale bar = 100 um for all images.

### Effects of AICAR treatment on autophagy following muscle injury

Given that our previously published work demonstrated that autophagy plays an important role in regeneration of muscle following injury^16^, we investigated various markers of autophagy as a potential mechanism for AICAR aiding in recovery of muscle function. Autophagy is a cellular recycling process that identifies, removes, and degrades damaged or dysfunctional cellular components via autophagosomes. Beclin1 (or Atg6) expression, an important marker of cellular autophagy which contributes to the formation of autophagosomes, was greater in KO mice compared to LM mice independent of injury or treatment (interaction P=0.039; Fig. 6a). Interestingly, AICAR treatment resulted in greater Beclin1 expression in the two-week post-injury limb of LM mice, but not KO mice (Fig. 6a).

**Figure 6:**
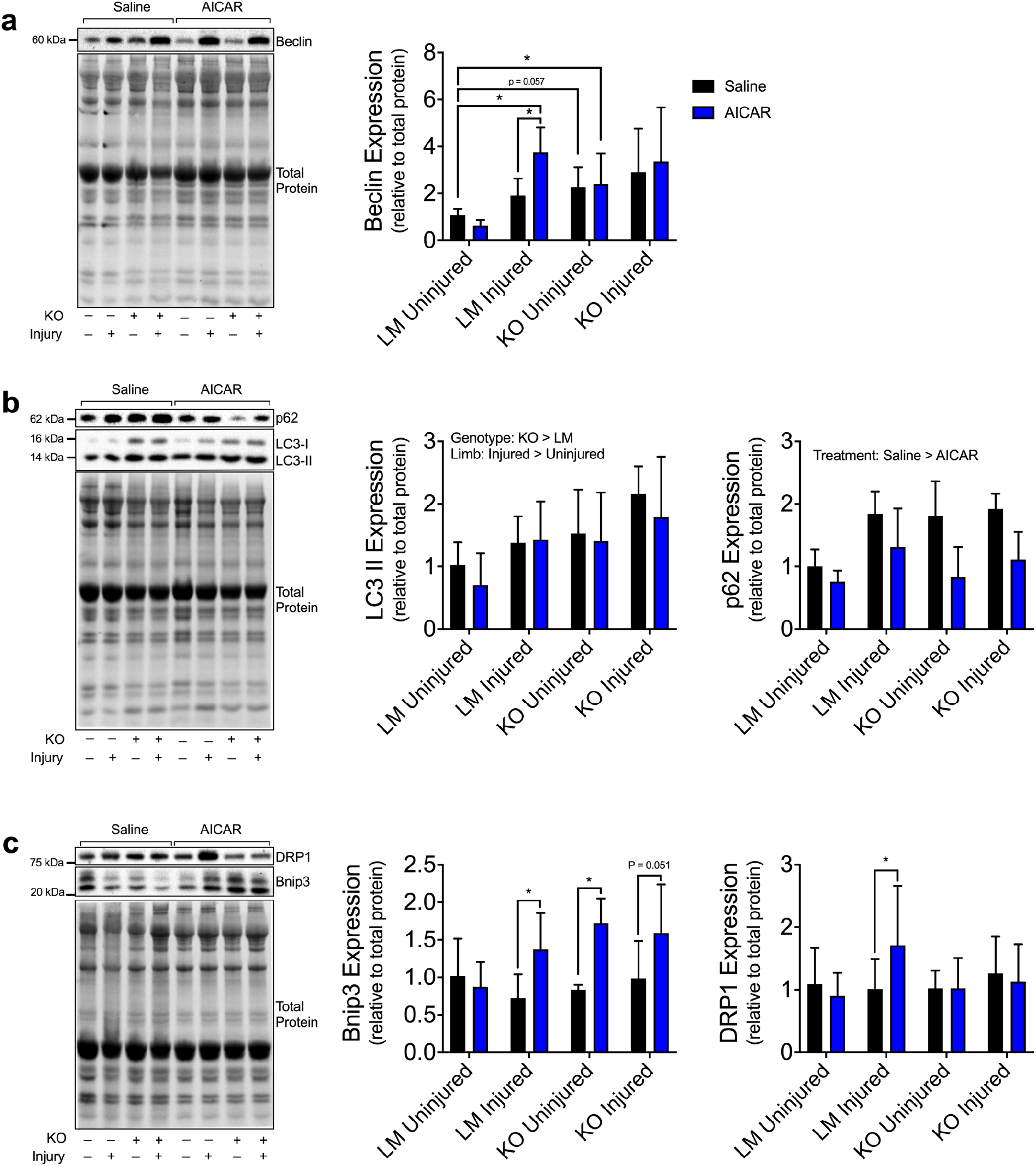
AICAR treatment aids in recovery of muscle strength following muscle injury. (a) Pre-cardiotoxin injury muscle strength from LM and KO mice treated with saline (LM n=13, KO n=9) or AICAR (LM n=13, KO n=7; two-way ANOVA, main effect of genotype P<0.001) (b) Muscle strength 14 days after cardiotoxin injury for LM and KO mice treated with saline (LM n=12, KO n=9) or AICAR (LM n=10, KO n=6; main effect of treatment P=0.006). Data are presented as means ± SD.

The final step of autophagy is the degradation of the autophagasome and the level of activity of this process can be characterized by measuring protein content levels of the conjugated form of microtubule-associated proteins 1A/1B light chain 3A (LC3-II), as well as the relative degradation of polyubiquitin-binding protein p62/SQSTM1 (p62). LC3-II content was elevated by ~52% in fukutin KO mice compared to LM mice (Fig. 6b; P<0.002) and by ~32% in the injured limb compared to uninjured limb (Fig. 6b; P<0.001), but these effects were independent of AICAR treatment. Interestingly, degradation of p62 was ~54% greater in AICAR-treated mice compared to untreated mice (Fig. 6b; P=0.031).

BCL2/adenovirus E1B 19 kDa protein-interacting protein 3 (Bnip3) and dynamin-related protein 1 (Drp1) are proteins involved in mitochondrial specific autophagy, known as mitophagy. Bnip3, which is specifically responsible for tagging mitochondria for degradation, was ~60-100% greater in the AICAR-treated LM injured, KO uninjured, and KO injured mice compared to corresponding untreated mice (Fig. 6c; limb x genotype x treatment interaction: P=0.016). Drp1 which facilitates removal of damaged mitochondria from the network was also elevated (~70%) in response AICAR treatment, but only in LM-injured AICAR-treated mice (Fig. 6c; limb x genotype x treatment interaction: P=0.037).

## Discussion

Disruptions in the function of αDG within the DGC will give rise to a collection of muscle myopathies termed secondary dystroglycanopathies that are characterized by early onset of muscle weakness and progressive loss of muscle function. Importantly, in order to develop targeted treatments for these myopathies, we must first understand how disruptions in the DGC may manifest in muscle pathophysiology. In this study, we reveal that fukutin deletion leads to reductions in mitochondrial function and content compared to LM control mice, which is likely the result of impaired mitochondrial biogenesis. These results point to the possibility of a link between muscle metabolic function and proper function of αDG meaning that the resulting metabolic disruptions stemming from loss of fukutin could be an important component of the disease pathology. Secondly, we demonstrate that the AMPK agonist AICAR potentially accelerates muscle recovery from injury, making the AMPK signaling pathway an intriguing option for future research involving treatments for dystroglycanopathies.

In recent years, it has become apparent that mitochondrial abnormalities are a common phenotype presenting in mice with various forms of DGC disruptions^13,18–20^. However, research involving mitochondria in the context of dystroglycanopathies has yet to elucidate whether the mitochondrial abnormalities are intrinsic manifestations or secondary consequences of disease pathology. Herein, we present data that suggests that mitochondrial defects are intrinsic to the disease pathology as KO mice have impaired PGC-1α transcription following acute exercise. This data suggests that poor mitochondrial quality in KO mice is likely stemming from modifications to the PGC-1α gene that is derived from altered DGC signaling. This idea is supported by the recent work performed by Pambianco and colleagues who demonstrated that DGC disruption via deletion of α-sarcoglycan leads to reduced acetylation and a condensed chromatin structure of the PGC-1α promoter^18^. This nuclear modification arising from a dysfunctional DGC resulted in diminished mitochondrial function due to reduced mitochondrial proteins that are under the regulatory control of PGC-1α. PGC-1α is likely not the only metabolic gene that affected by disruptions in the DGC, as gene expression profiling performed on human tissue revealed that 26 nuclear encoded mitochondrial genes were downregulated in dystrophin and α-SG deficient patients^21^. The downregulation of these metabolic genes could potentially be the result of overactive Ca^2+^ signaling in the muscle cells creating a negative feedback loop that drives down the metabolic gene expression. Together these data suggest that mitochondrial defects are linked to altered DGC function, but more research is necessary to explore whether these defects are directly linked to or simply accompany the DGC disruptions. Nevertheless, this data does provide an potential interesting avenue for future research regarding treatments strategies for secondary dystroglycanopathies.

Activation of the AMPK signaling pathway is one of the primary mechanisms underlying mitochondrial biogenesis, and AICAR, being an AMPK agonist, is a drug that has been shown to stimulate the mitochondrial biogenesis process. In this study, we attempted to exploit the benefits of AICAR on mitochondrial biogenesis to rescue the mitochondrial defects observed in KO mice. However, AICAR had no beneficial effects on mitochondrial content or function, which could potentially be due to faulty PGC-1α transcription given that AICAR stimulates mitochondrial biogenesis through AMPK-induced activation of PGC-1α. Minimal effects of AICAR on mitochondria have also been reported by Pauly and colleagues^13^ who treated dystrophin-deficient *mdx* mice with AICAR for 4 weeks and found no changes in mitochondrial content or respiratory function. Irrespective of the reason for AICAR having minimal effects on muscle mitochondria, this data informs future research to employ other drugs to target alternate pathways in order to correct mitochondrial defects in fukutin KO mice.

Our group has previously published a report with histological evidence suggesting that KO mice have abnormal muscle regeneration following severe muscle injury^8^. Thus, to determine whether the histological alterations in KO mice translated to functional deficits in recovery and to see if AICAR could help correct any abnormal recovery, we measured muscle function in LM and KO mice with and without AICAR treatment following an injury. Interestingly, despite our previously published data, fukutin KO mice did not have any signs of delayed regeneration at 2 weeks following the injury. This is not to say that a delay in functional recovery would never happen in fukutin KO mice, as one injury may not be sufficient to observe defects in functional recovery. Perhaps, repeated cardiotoxin injuries would have revealed a regeneration deficit that only becomes apparent after chronic repeated injuries. Our results demonstrate that KO mice have the ability to undergo normal functional regeneration following a severe injury, but it remains to be seen whether further injuries or aging would have an effect on this finding.

While ACIAR treatment did not have any beneficial effects on mitochondria in fukutin KO mice, it did have a beneficial effect on recovery of muscle function following muscle injury in both LM and KO mice. Initially, we hypothesized that AICAR treatment would be beneficial for KO mice after injury because it would enhance mitochondrial maintenance but given the lack of effect of AICAR on mitochondria, it remains to be seen how AICAR would lead to improved recovery after injury. AICAR is known to stimulate autophagy, and we have previously reported that enhancing autophagy prior to an injury can lead to accelerated recovery of muscle function^16^. In this study, we found that AICAR treatment did indeed augment autophagy in both LM and KO mice, which is a possible explanation for AICAR’s effect on recovery of muscle function. Alternatively, AICAR has also been shown to induce satellite cell proliferation^22^, which may be a mechanism for enhancing recovery following injury^23^. Our findings reveal the potentially promising application of AICAR as a way to speed functional recovery of muscle after injury, and future research should seek to further explore AICAR’s mechanism of action in order to optimize treatment efficacy.

## Acknowledgements

This work was funded, in part, by the Muscular Dystrophy Foundation Research Grant #480351(Beedle).

